# PU.1 and TGF-β signaling transactivate CD103 expression in mast cells and dendritic cells: Opposite roles of GATA2 in the expression of mucosal mast cell-specific genes

**DOI:** 10.1101/2025.02.10.637371

**Authors:** Kenta Ishii, Kazuki Nagata, Niya Yamashita, Yuki Yamazaki, Yuta Akimoto, Weiting Zhao, Mariko Inoue, Naoto Ito, Kazumi Kasakura, Chiharu Nishiyama

**Affiliations:** Department of Biological Science and Technology, Faculty of Advanced Engineering, Tokyo University of Science, 6-3-1 Niijuku, Katsushika-ku, Tokyo 125-8585, Japan

## Abstract

Mucosal mast cells (MMCs) are distinguished from connective tissue MCs by the specific expression of integrin CD103 (αE/β7) and MC proteases Mcpt1 and Mcpt2. Although the expression of the *Mcpt1* and *Mcpt2* genes is cooperatively regulated by the transcription factor GATA2 and TGF-β signaling in MMCs, the transcriptional mechanism of CD103 expression remains unknown. Here, we found that surface CD103 and *Itgae* mRNA levels were significantly increased by the knockdown (KD) of *Gata2* in bone marrow-derived MCs (BMMCs), which was accelerated by a TGF-β stimulation. Since the mRNA levels of *Spi1* (encoding PU.1) were increased in *Gata2* KD BMMCs, we examined the effects of PU.1 on CD103 expression. As expected, CD103 levels on BMMCs were significantly decreased by *Spi1* KD and increased by *Spi1* overexpression. *Spi1* KD suppressed *Itgae* expression even in the presence of TGF-β in BMMCs and peritoneal MCs, whereas *Gata2* KD amplified the TGF-β-induced increase in *Itgae* expression. The amount of PU.1 binding to the *cis*-element in the *Itgae* gene was significantly and moderately increased by *Gata2* KD and the TGF-β stimulation, respectively. Since PU.1 is an essential transcription factor for dendritic cells (DCs), we examined the role of PU.1 in CD103 expression on DCs. The KD experiment using BMDCs showed a significant decrease in CD103 levels in *Spi1* siRNA-transfected BMDCs.

We concluded that PU.1 affected CD103 expression on MMCs and DCs by transactivating the *Itgae* gene, and also that GATA2, which positively regulated the MMC-specific expression of Mcpt1 and 2, inhibited CD103 expression by repressing PU.1.

## Introduction

Mast cells (MCs), which play an important role in the pathogenesis of allergic diseases and defenses against parasitic infection, are classified into two subtypes: mucosal MCs (MMCs) and connective tissue MCs (CTMCs). MMCs and CTMCs are distinguished by their specific gene expression profiles. In mice, MMCs specifically express mast cell protease (Mcpt)1, Mcpt2, and integrin CD103, whereas CTMCs express Mcpt4 and 5. MMCs and CTMCs acquire this cell type-specific gene expression profile during the tissue environment-dependent maturation of bone marrow-derived immature MCs at mucosal and connective tissues, respectively. Therefore, the molecular mechanisms of transcriptional regulation, which controls the commitment between MMCs and CTMCs, need to be clarified in order to obtain a more detailed understanding of differences in the development and function of MMCs and CTMCs. We previously investigated transcription factors regulating the expression of the *Mcpt1* and *Mcpt2* genes and identified the hematopoietic cell-specific transcription factor GATA2 [1]. Although we revealed the essential role of GATA2 in the expression of the *Mcpt1* and *Mcpt2* genes, which was driven through tandemly repeated *cis*-elements in the proximal region of these genes in a cooperative manner with the TGF-β-Smad axis, the mechanisms by which the fate of MMCs and CTMCs is decided remains unclear because GATA2 is a positive regulator of the development of all MCs, basophils, and their common progenitor. To investigate MMC-specific gene regulation, we herein examined the expression mechanism of CD103 in MCs.

CD103, a heterodimeric cell surface molecule composed of αE and β7 integrins, is mainly expressed on hematopoietic cells, including T cells and dendritic cells (DCs), and is involved in the localization of immune cells to peripheral tissues, particularly the intestinal mucosa, by interacting with E-cadherin expressed on epithelial cells. Based on our previous study showing that GATA2 was an essential positive regulator of Mcpt1 and Mcpt2 expression in MMCs [1], we herein examined the effects of the knockdown (KD) of *Gata2* on CD103 expression in MMCs. CD103 expression was significantly up-regulated by *Gata2* KD due to an increase in the mRNA level of *Itgae* (encoding αE), which suggested opposite roles for GATA2 in MMC-specific gene expression, i.e., as a suppressor of *Itgae* gene and activator of *Mcpt1* and *2* genes. We then focused on the transcription factor PU.1 because it is in a negative crosstalk relationship with GATA2 and/or GATA1 in various hematopoietic lineages [2–9], and also plays a cooperative role with GATA1/2 in MCs [10–12]. We found that the function and expression of PU.1 was up-regulated in *Gata2* KD MCs, and the KD of *Spi1* (encoding PU.1) significantly reduced CD103 expression not only on MCs, but also on DCs. In addition, we investigated the involvement of other transcriptional mechanisms, including TGF-β signaling and epigenetic regulation, on the expression of CD103.

## Materials and Methods

### Mice and Cells

BMMCs and BMDCs were generated from the BM cells of male C57BL/6J mice (Japan SLC, Hamamatsu, Japan), which were maintained under specific pathogen-free-conditions, by their maintenance in the presence of 5 μg/mL recombinant mouse IL-3 (#575508, BioLegend) [1, 13], and 100 ng/mL recombinant mouse Flt3L (#575508, BioLegend), respectively. Peritoneal MCs were prepared from male C57BL/6J mice as previously described [14].

All experiments using mice were performed following the guidelines of the Institutional Review Board of Tokyo University of Science, and the present study was approved by the Animal Care and Use Committees of Tokyo University of Science: K22005, K21004, K20005, K19006, K18006, K17009, K17012, K16007, and K16010.

### Quantification of mRNAs

Total RNA was extracted from cells using the ReliaPrep RNA Cell Miniprep System (#Z6012), or RNAzol RT Reagent (#RN190, Cosmo Bio). Complementary DNA was synthesized from total RNA using ReverTraAce qPCR RT Master Mix (#FSQ-201, TOYOBO). Quantitative PCR was performed by the StepOne Real-Time PCR System (Applied Biosystems) with THUNDERBIRD probe qPCR Mix (#QPS-101, TOYOBO) or THUNDERBIRD SYBR qPCR Mix (#QPS-201, TOYOBO) using the following primers.

*Gata2*: Mm00492300_m1 (Applied Biosystems) or a primer set with forward; 5’-CCCAAGCGGAGGCTGTCT-3’, and reverse; 5’-TGTCGTCTGACAATTTGCACAA-3’.

*Spi1*: Mm00488142_m1 (Applied Biosystems) or a primer set with forward; 5’-ATGTTACAGGCGTGCAAAATGG-3’, and reverse; 5’-TGATCGCTATGGCTTTCTCCA-3’.

*Irf4*: a primer set with forward; 5’-CCCCATTGAGCCAAGCATAA-3’, and reverse; 5’-GCAGCCGGCAGTCTGAGA-3’.

*Cebpa*: Mm00514283_s1 (Applied Biosystems) or a primer set with forward; 5’-CGCAAGAGCCGAGATAAAGC-3’, and reverse; 5’-CGCAGGCGGTCATTGTC-3’.

*Itgae*: forward; 5’-TAGGCAAGATGGATGTAAAACTGTGT-3’, and reverse; 5’-CTCTCTAAGGCCTGGCTCAGAA-3’.

*Itgb7*: forward; 5’-CAAGTCACCATGTGAGCAG-3’, and reverse; 5’-GTCAAGGTCACATTCACGTC-3’.

*Mcpt1*: forward; 5’-AAGTTCCACAAAGTTAAAAACAGCATAC-3’, reverse; 5’-GTGAATCCCCATAAGATACAATACCAT-3’.

*Mcpt2*: forward; 5’-AAAGTTTCAGTACCTTTCGGG-3’, and reverse; 5’-CATCCACATCAGAATTCAACTCT-3’.

*Mcpt4*: forward; 5’-GTTCACCCAAAGTACAACTTCTA-3’, and reverse; 5’-ATTCACAGAGGGAGTCTCTTTG-3’.

The mRNA levels of target genes were evaluated as a ratio against that of *Gapdh* (4352339E or a primer set of forward; 5’-ACGTGCCGCCTGGAGAA-3’ and reverse; 5’-GATGCCTGCTTCACCACCTT-3’) as the housekeeping gene by calculating cycle threshold values.

### Flow cytometric analysis

Cells were preincubated with 1 μg/mL Fc block and 1 μg/mL DAPI on ice for 5 min and were stained with PE-labeled anti-CD103 Ab (clone 2E7, Myltenyi Biotec, 1:50 or BioLegend, 1:300), APC-labeled anti-CD117/c-kit Ab (clone ACK2, BioLegend, 1:500), PE/Cy7- or FITC-labeled anti-FcεRIα Ab (clone 2E7, BioLegend, 1:500), PE/Cy7-labeled anti-mouse CD11c (clone N418, BioLegend, 1:1000), APC-labeled anti-mouse PDCA1 (clone 927, BioLegend, 1:300), FITC-labeled anti-mouse/human CD45R/B220 (clone RA3-6B2, BioLegend, 1:300), APC/Cy7-labeled anti-human/mouse CD11b (clone M1/70, BioLegend, 1:2000), or PE Hamster IgG2k (BD BioSciences) as an isotype control of anti-CD103 Ab, on ice for 15 min. After washing, stained cells were detected using a FACSLyric Analyzer (BD Pharmingen) or a MACS Quant Analyzer (Miltenyi Biotec). Data were processed with FlowJo software (Tomy Digital Biology).

### Transfection of siRNA

We introduced siRNA into MCs with a Neon 100 μl kit (Thermo Fisher Scientific) using a Neon Transfection System (Thermo Fisher Scientific) and into BMDCs with the mouse dendritic cell nucleofector kit (Lonza, Basel, Switzerland) by Nucleofector II (Lonza).

The following siRNAs purchased from Invitrogen (Carlsbad, CA) were used for KD experiments: *Gata2* siRNA (MSS204585), *Spi1* siRNA (MSS247676), *Irf4* siRNA (MSS273622), *Smad3* siRNA (MSS20642), *Smad4* siRNA (MSS20643), and their GC content-matched appropriate controls from the Stealth RNAi siRNA Negative Control Kit (#12935100).

### Western blot analysis

Western blot analyses were performed as previously described [1, 15] with anti-GATA2 Ab (CG2-96, sc-267, and H-116, sc-9008, Santa Cruz Biotechnology), anti-PU.1 Ab (C-3, sc-390405, and D19, sc-5949, Santa Cruz Biotechnology), anti-β-actin Ab (AC-15, A5441, Sigma-Aldrich), and anti-GAPDH (14C-10, 2118, Cell Signaling Technology).

### ChIP assay

ChIP assays were performed as previously described [1, 13, 15]. Anti-PU.1 Ab (T21, Santa Cruz Biotechnology), anti-acetyl histone H4 Ab (06-866, Millipore), or control rabbit IgG (02-6102, Invitrogen) was used. The nucleotide sequences of the forward (F) and reverse (R) primers to detect chromosomal DNA were as follows:

*Itgae* region1,

F; 5’-CCCCTGGTCCTGTTGCTAAG-3’, R; 5’-CTGGCGGGCAGTGACTTC-3’

*Itgae* region2,

F; 5’-CCTGGCCTAGACCCCAGAAT-3’, R; 5’-TCCGAGTATCTGGGAGGTTGA-3’

*Itgae* region3,

F;5’-CACCCCGATGCCATGTAGTC-3’,

R;5’-TTACCATGGTCACAGAATGAGTCA-3’

*Itgae* region4,

F; 5’-GGGCAGGAAATGCACCAA-3’, R; 5’-CGGCTATCAGCCTGTTCCTT-3’

*Itgae* region5,

F; 5’-TGCTCTTCCCCTCAGCCTTT-3’, R; 5’-CATCTGGGTGGATCCCTGAA-3’

*Itgae* region6,

F;5’-CCTGTGCTGCAGAGAGGAAGT-3’, R;5’-GAGGACCAGACAGCTCAGCAT-3’

*Itgae* promoter,

F; 5’-TGCCAAAGACACTCAGAACGA-3’, R; 5’-CACCATCTCATGCACGAGTGA-3’

*Mcpt2* promoter,

F; 5’-TGCCAAAGACACTCAGAACGA-3’, R; 5’-CACCATCTCATGCACGAGTGA-3’

### Overexpression of *Spi1*

We used an expression plasmid of *Spi1*, pIRES2-AcGFP-3FL-rPU.1 [16], generated in our previous study by the insertion of 3×Flag-tagged rat *Spi1* cDNA into pIRES2-AcGFP1 (Takara Bio). The transfection of BMMCs with pIRES2-AcGFP-3FL-rPU.1 or pIRES2-AcGFP1 (control) was performed using the Neon Transfection System as well as siRNA transfection.

### Statistical analysis

A two-tailed Student’s t-test was used to compare two samples. To compare more than three samples, a one-way ANOVA followed by Tukey’s multiple comparison test or Dunnett’s multiple comparison test was performed. *p*-Values <0.05 were considered to be significant.

## Results

### The TGF-β stimulation induced the surface expression of CD103 accompanied by an increase in *Itgae* mRNA levels in MCs

TGF-β accelerates the development of MMCs specifically expressing Mcpt1, Mcpt2, and CD103, which are not observed in CTMCs [17]. Although we have previously revealed that GATA2 and Smads cooperatively transactivated the *Mcpt1* and *Mcpt2* genes by forming a complex in MCs [1], the molecular mechanisms by which the MMC-specific expression of CD103 is induced remain unclear.

Therefore, we examined the effects of TGF-β on the expression of MMC-related genes in BMMCs. When BMMCs were treated with 1 ng/mL TGF-β for 24 or 48 h, the apparent surface expression of CD103 was induced (**Fig. 1a**). Quantitative PCR revealed that the mRNA levels of *Itgae* but not *Itgb7,* were significantly increased in TGF-β-treated MCs (**Fig. 1b**). These results suggest that the BMMCs generated under our experimental conditions (IL-3 supplementation) possessed the potential for the high expression of the *Itgae* gene and then CD103, whereas an additional supplementation with SCF was required for the TGF-β-mediated induction of *Itgae* expression in IL-3-induced BMMCs in a previous study [17]. A time-course analysis showed that after the TGF-β stimulation, *Itgae* mRNA levels increased by approximately 5-fold at 4 h, 8-fold at 6 h, and were sustained at least until 48 h following an 80-fold increase at 24 h (**Fig. 1c**). The involvement of Smad3 and Smad4 was confirmed by a KD experiment using siRNA against *Smad3* and *Smad4*, in which the TGF-β-induced increase in *Itgae* mRNA levels was significantly suppressed by *Smad3* KD or *Smad4* KD (**Fig. 1d**). The TGF-β stimulation also significantly increased the mRNA levels of *Mcpt1* and *Mcpt2* (**Fig. 1b and c**).

**Fig. 1.**
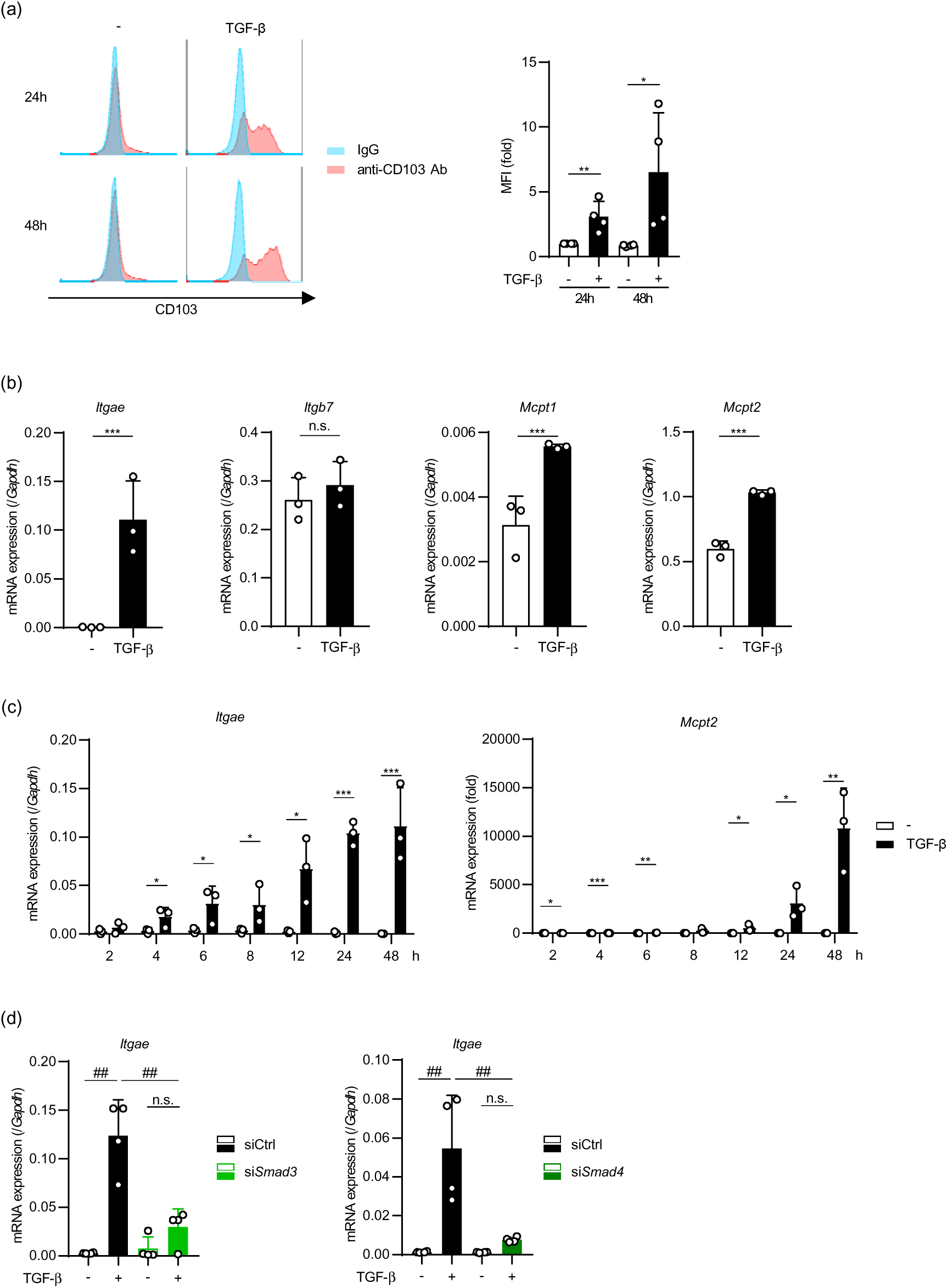
Effects of the TGF-β stimulation on the expression of MMC-related genes in BMMCs. **a.** Cell surface levels of CD103 on BMMCs. BMMCs were generated from C57BL/6J male BM cells through cultivation in the presence of 5 ng/mL recombinant mouse IL-3 for 6-8 weeks. Left; Typical histograms of staining with anti-CD103 Ab or its control of the FcεRI^+^/c-kit^+^ population (>95% of total BMMCs) in BMMCs, which were treated with 1 ng/mL recombinant mouse TGF-β for 24 h or 48 h. Right; Mean fluorescence intensities (MFIs) obtained from four independent experiments. The MFI ratio obtained with the following calculation are shown: (MFI stained with anti-CD103 Ab)/(MFI stained with the isotype control). In all experiments in the present study, independent experiments were performed on different days using BMMCs prepared independently from different individuals. **b.** mRNA levels of *Itgae*, *Itgb7*, *Mcpt1*, *Mcpt2*, and *Mcpt4* in BMMCs incubated in the presence or absence of 1 ng/mL TGF-β for 48 h. **c.** mRNA levels of *Itgae* and *Mcpt2* in BMMCs after an incubation with or without 1 ng/mL TGF-β for the indicated times. **d.** Effects of the KD of Smads on the TGF-β-induced increase in *Itgae* mRNA levels. Fourty-eight hours after electroporation for siRNA transfection, BMMCs were stimulated with 1 ng/mL TGF-β for an additional 6 h. Data represent the mean ± SEM of independent experiments. Dunnett’s multiple comparison test (**a**, **d**) and the two-tailed paired *t*-test (**b**) were used for statistical analyses. *, *p* < 0.05; ***, *p* <0.005; n.s., not significant.

### *Gata2* KD increased surface CD103 and *Itgae* mRNA levels, but decreased *Mcpt2* mRNA levels in MCs

GATA2, which plays an important role in the development and gene expression of MCs and basophils, has been identified as a transactivator of the *Mcpt1* and *Mcpt2* genes in MMCs [1]. To elucidate the role of GATA2 in the expression of CD103 in MCs, we introduced *Gata2* siRNA into BMMCs. GATA2 protein and *Gata2* and *Mcpt2* mRNA levels were significantly decreased in *Gata2*-KD BMMCs, which is consistent with previous findings [1], whereas *Itgae* and *Itgb7* mRNA levels were markedly and significantly increased, respectively, by *Gata2* KD (**Fig. 2a and b**). The surface expression of CD103 on MCs was also strongly induced by *Gata2* KD (**Fig. 2c**). In contrast to the TGF-β-induced transactivation of *Mcpt2*, which was largely abolished in *Gata2*-KD MCs, that of *Itgae* was significantly amplified by *Gata2* KD (**Fig. 2d**). The synergistic enhancing effects of *Gata2* KD on the TGF-β-induced increase in *Itgae* transcripts were reflected in the surface protein level of CD103 on MCs (**Fig. 2e**).

**Fig. 2.**
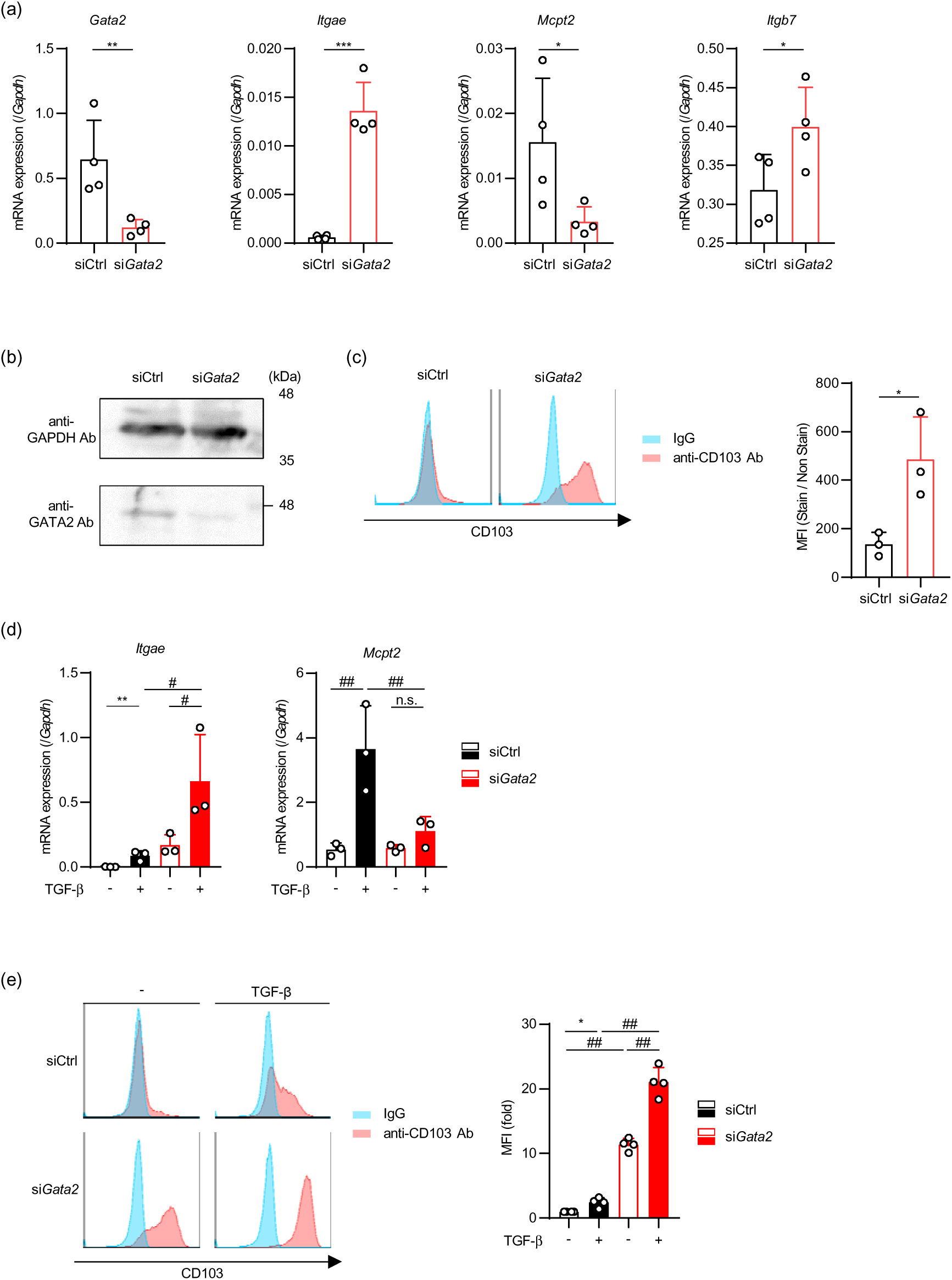
Effects of *Gata2* KD on the expression of MMC-related genes in BMMCs. **a.** mRNA levels of *Gata2*, *Itgae*, *Itgb7*, and *Mcpt2* in *Gata2* siRNA-transfected BMMCs. **b.** Western blotting profiles of *Gata2* KD and control BMMCs. Whole cell lysates containing 20 mg of protein were loaded onto each lane. After the detection of GATA2 proteins, the stripped membrane was re-probed with anti-β-actin Ab. **c.** Cell surface CD103 levels on *Gata2* siRNA-transfected BMMCs. **d.** mRNA levels of *Itgae* and *Mcpt2* in *Gata2* siRNA-transfected and TGF-β-treated BMMCs. **e.** Cell surface CD103 levels on *Gata2* siRNA-transfected and TGF-β-treated BMMCs. BMMCs transfected with *Gata2* siRNA or its control siRNA were harvested 48 h after transfection (**a**-**c**), *Gata2* siRNA transfectants or its controls were collected 48 h after electroporation and replaced in medium with or without TGF-β for an additional 48-h incubation (**d**, **e**). Data represent the mean ± SEM of independent experiments. Dunnett’s multiple comparison test (**d**, **e**) and the two-tailed paired *t*-test (**a, c**) were used for statistical analyses. *, *p* < 0.05; **, *p* <0.01; n.s., not significant.

These results indicate that GATA2 has opposite roles in MMC-related genes; namely, as a positive regulator of the *Mcpt1* and *Mcpt2* genes and as a negative regulator of the *Itgae* and *Itgb7* genes.

### Effects of *Gata2* KD on *Spi1* mRNA levels, and of *Spi1* KD on surface CD103 and *Itgae* mRNA levels in MCs

GATA2 and/or GATA1 suppress the function and expression of PU.1 through the development of erythroid/megakaryocyte lineages from hematopoietic stem cells, whereas GATA2(1) and PU.1 cooperatively regulate the expression of MC-specific genes [10, 11]. To investigate whether the expression of the *Spi1* gene in MCs is regulated by GATA2, we performed quantitative PCR on mRNAs in BMMCs and found that *Spi1* mRNA levels were significantly increased by *Gata2* KD (**Fig. 3a**). In addition, we revealed that the mRNA level of *Irf4* was low and not affected by *Gata2* KD in BMMCs (**Fig. 3a**), while IRF4 is a partner molecule of PU.1 and is reportedly required for CD103^+^ DC survival *in vivo* [18]. *Spi1* KD, but not *Irf4* KD, significantly decreased *Itgae* mRNA levels (**Fig. 3b**), suggesting that *Gata2* KD up-retulated the expression of *Spi1*, which subsequently transactivated the *Itgae* gene in an IRF4-independent manner. We examined *Spi1* KD BMMCs and showed the mRNA levels of *Itgae* and *Itgb7* decreased and those of *Mcpt2* and *Gata2* increased (**Fig. 3c**) under the condition that *Spi1*KD effectively reduced PU.1 protein levels (**Fig. 3d**).

**Fig. 3.**
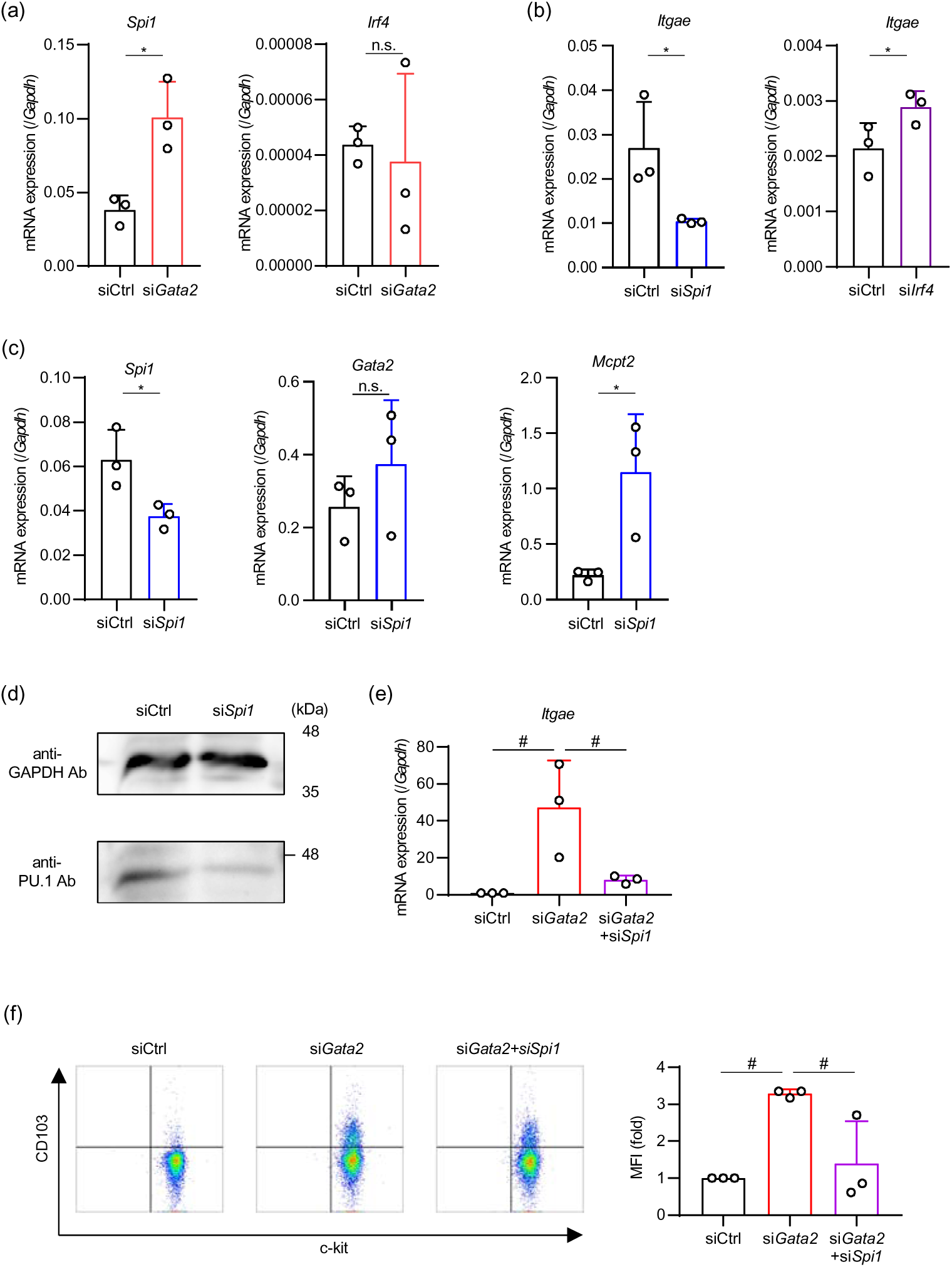
Role of PU.1 in the *Gata2* KD-induced up-regulation of CD103 on BMMCs. **a.** mRNA levels of *Spi1* and *Irf4* in *Gata2* KD BMMCs. **b.** *Itgae* mRNA levels in *Spi1* or *Irf4* KD BMMCs. **c.** mRNA levels of MMC-related molecules in *Spi1* KD BMMCs. **d.** Western blotting profile of PU.1 protein levels in *Spi1* KD BMMCs. **e.** and **f.** Effects of *Spi1* KD on the *Gata2* KD-induced up-regulation of *Itgae* mRNA (**e**) and of cell surface CD103 (**f**). A typical dot-plot profile of staining with anti-CD103 Ab and anti-c-kit (left in **f**) and MFIs of surface CD103 obtained from three independent experiments (right in **f**). Data represent the mean ± SEM of independent experiments. Dunnett’s multiple comparison test (**e**, **f**) and the two-tailed paired *t*-test (**a**, **b**, **c**, **d**) were used for statistical analyses. *, *p* < 0.05; n.s., not significant.

To investigate the involvement of PU.1 in the *Gata2*-KD-induced up-regulation of CD103, we performed the doble KD of *Gata2* and *Spi1*. *Gata2*-KD-mediated increases in the expression of *Itgae* mRNA (**Fig. 3e**) and CD103 (**Fig. 3f**) were significantly abrogated by the additional KD of *Spi1*.

These results suggest that *Gata2* KD up-regulated CD103 by enhancing the PU.1-dependent transcription of the *Itgae* and *Itgb7* genes.

### The amount of PU.1 binding to the *Itgae* gene in MCs was increased by *Gata2* KD

To evaluate the effects of *Gata2* KD on PU.1 function related to the expression of the *Itgae* gene in MCs, we conducted a ChIP assay. Based on the ChIP-Atlas database, we selected 6 regions as candidates for PU.1-binding sites around the *Itgae* gene (**Fig. 4a**). Region3 and region4 were identified in a ChIP assay using anti-PU.1 Ab as the regions most strongly binding with PU.1 in MCs (**Fig. 4b**). We then performed a ChIP assay to measure the amount of PU.1 binding to the *Itgae* gene targeting region3 in *Gata2*-KD BMMCs. As shown in **Fig. 4c**, the binding degree of PU.1 to this region in the *Itgae* gene was significantly increased by *Gata2* KD.

**Fig. 4.**
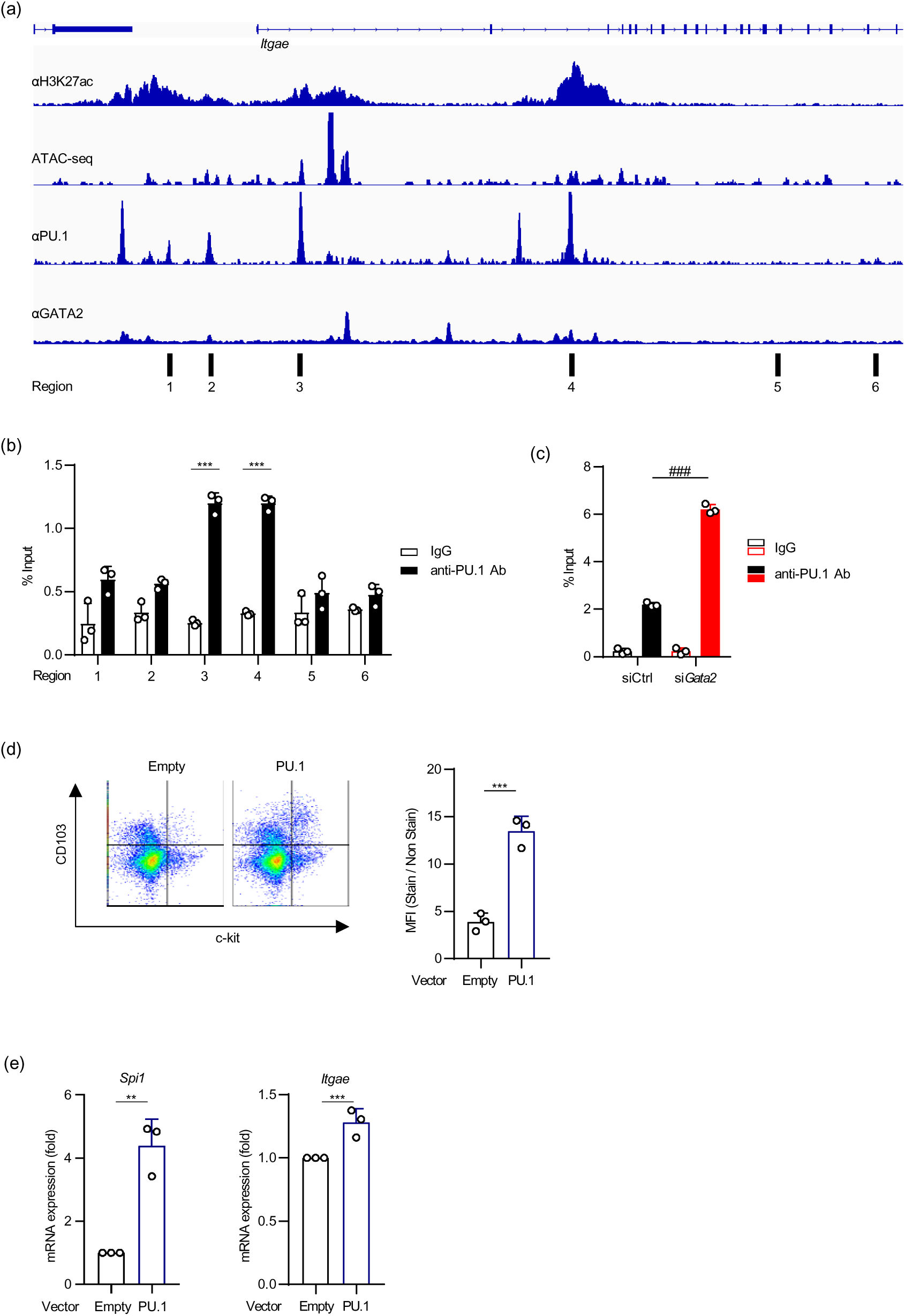
Effects of *Gata2* KD on PU.1 occupancy in the *Itgae* gene, and of PU.1 overexpression on CD103 levels. **a.** The profiles of H3K27 acetylation, ATAC-seq, and PU.1-binding around the *Itgae* gene in MCs. H3K27 acetylation (https://chip-atlas.dbcls.jp/data/mm9/eachData/bw/SRX1456419), ATAC-seq (https://chip-atlas.dbcls.jp/data/mm9/eachData/bw/SRX7750160), PU.1-binding (https://chip-atlas.dbcls.jp/data/mm9/eachData/bw/SRX5026215), and GATA2-binding (https://chip-atlas.dbcls.jp/data/mm9/eachData/bw/SRX9030838) were obtained via the ChIP-Atlas analysis (http://chip-atlas.org). **b.** Amount of PU.1 binding to 6 selected regions around the *Itgae* gene in BMMCs. **c.** Effects of *Gata2* KD on the amount of PU.1 binding to region3 in the *Itgae* gene. **d.** Surface CD103 level on *Spi1*-overexpressed BMMCs. A typical dot plot profile detected with GFP and anti-CD103 Ab (left) and MFIs of surface CD103 obtained from three independent experiments (right). Empty; BMMCs transfected with mock vector (pIRES2-AcGFP), PU.1; BMMCs transfected with pIRES2-AcGFP-3FL-rPU.1. **e.** mRNA level of *Itgae* in total BMMCs subjected to electroporation with pIRES2-AcGFP (empty) or pIRES2-AcGFP-3FL-rPU.1 (PU.1). Data represent the mean ± SEM of independent experiments. Dunnett’s multiple comparison test (**c**) and the two-tailed paired *t*-test (**b**, **d**, **e**) were used for statistical analyses. *, *p* < 0.05; n.s., not significant.

### Enforced expression of *Spi1* in MCs increased CD103^+^ cells

We exogenously overexpressed *Spi1* in BMMCs using a transient expression plasmid carrying IRES-GFP. Transfection of BMMCs with the *Spi1* expression plasmid resulted in the emergence of CD103^+^/GFP^+^ cells, which were absent in the mock transfectants (**Fig. 4d**). A significant increase in *Itgae* mRNA levels following the overexpression of *Spi1* was detected in a qPCR analysis of total BMMCs (without the isolation of GFP^+^ cells) (**Fig. 4e**), even though transfection efficiency was <10%. Quantitative PCR using a primer set specific for mouse *Spi1* mRNA showed that the endogenous transcription of *Spi1* (mouse *Spi1*) was significantly up-regulated by exogenous *Spi1* (rat *Spi1*) (**Fig. 4e**).

### The TGF-β stimulation enhanced the recruitment of PU.1 to the *Itgae* gene in MCs

In addition to the enhanced acetylation of histone H4 on the *Mcpt2* gene in TGF-β-treated BMMCs as observed in our previous study [1], the acetylation of H4 on the *Itgae* gene was significantly increased by the TGF-β stimulation (**Fig. 5a**). The TGF-β stimulation increased the amount of PU.1 binding to the *Itgae* gene (**Fig. 5b**), but did not affect the expression of PU.1 or GATA2 at the mRNA (**Fig. 5c**) or protein (**Fig. 5d**) levels. *Spi1* KD significantly suppressed the TGF-β-induced increase of CD103 expression on MCs (**Fig. 5e**), and this was accompanied by a decrease in the mRNA levels of *Itgae* and *Itgb7,* even in the presence of TGF-β, demonstrating the critical role of PU.1 in the expression of CD103 in MCs (**Fig. 5f**). Furthermore, the TGF-β-induced increase in *Mcpt2* transcription was abrogated by *Spi1* KD, which suggested the involvement of PU.1 in the TGF-β-smad-axis in MCs (**Fig. 5f**).

**Fig. 5.**
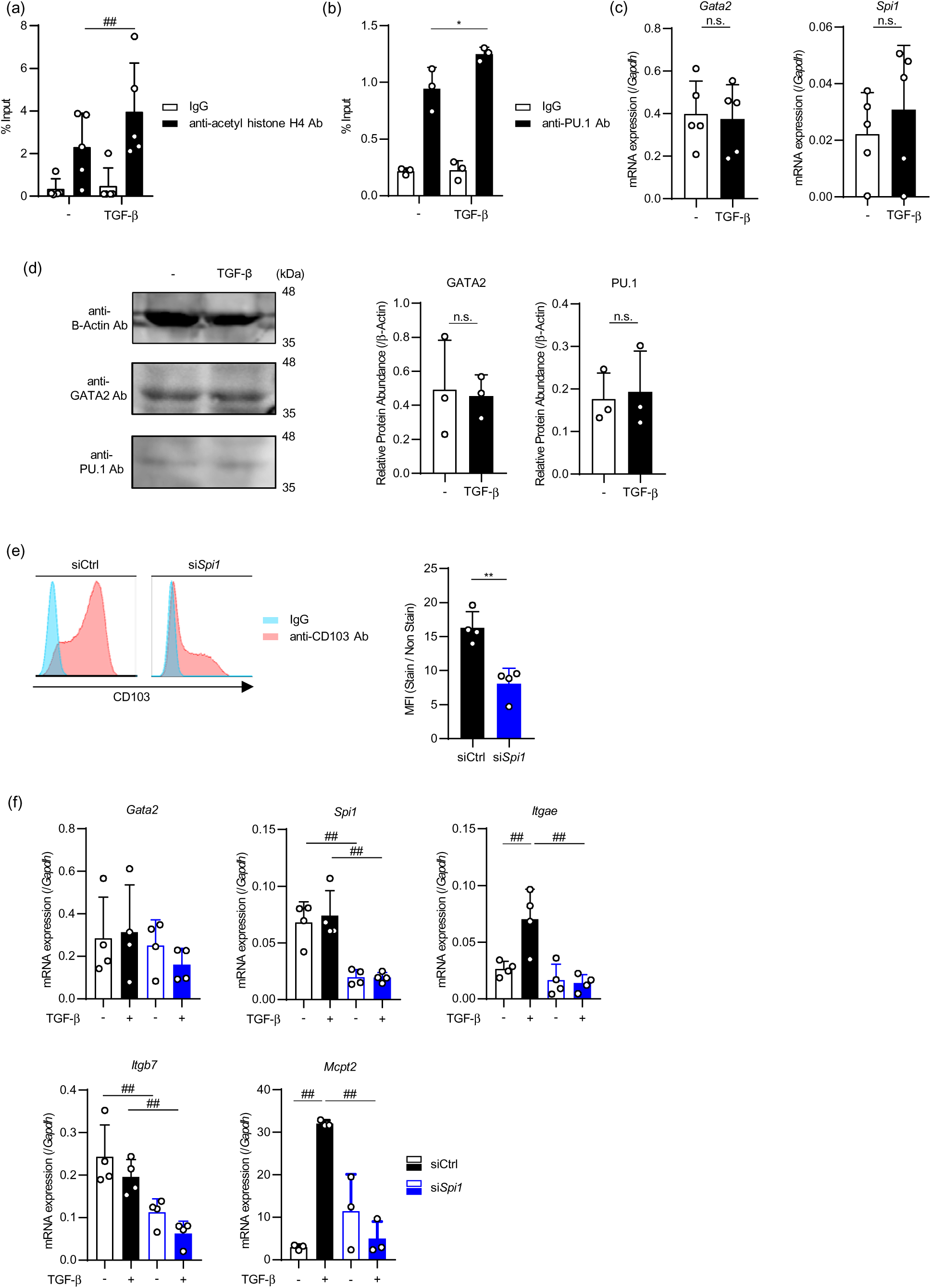
Effects of the TGF-β stimulation on PU.1 binding to the *Itgae* gene in BMMCs. **a.** ChIP assay data showing acetyl histone H4 on the *Itgae* and *Mcpt2* genes in TGF-β-treated BMMCs. **b.** ChIP assay data showing PU.1 binding to the region3 in the *Itgae* gene in TGF-β-treated BMMCs. **c.** and **d.** mRNA levels of *Spi1* and *Gata2* (**c**) and protein levels of PU.1 and GATA2 (**d**) in TGF-β-treated BMMCs. BMMCs were incubated in the presence or absence of TGF-β for 48 h (**a**-**d**). **e**. Surface CD103 levels on *Spi1* siRNA-transfected and TGF-β-treated BMMCs. **f.** mRNA levels of MMC-related genes in *Spi1* siRNA-transfected and TGF-β-treated BMMCs. BMMCs transfected with *Spi1* siRNA or its control were stimulated with TGF-β with the same time schedule as that in Fig. 2e. Data represent the mean ± SEM of independent experiments. Dunnett’s multiple comparison test (**b, e**) and the two-tailed paired *t*-test (**c**, **d**) were used for statistical analyses. *, *p* < 0.05; n.s., not significant.

### Effects of TGF-**β** signaling and the KD of *Gata2* or *Spi1* on peritoneal MCs

We investigated the involvement of TGF-β and PU.1 in the expression of the *Itgae* and *Itgb7* genes and CD103 using peritoneal MCs that were maintained with supplementation with 5 ng/mL SCF for 14 days for their survival. As observed in BMMCs, the TGF-β stimulation increased the mRNA levels of *Itgae* and *Mcpt2*, without affecting those of *Spi1*, *Gata2*, and *Itgb7* in peritoneal MCs (**Fig. 6a**). The cell surface expression of CD103 was detected on TGF-β-treated peritoneal MCs (**Fig. 6b**). It is important to note that *Gata2* KD significantly increased *Itgae* mRNA levels and amplified the TGF-β-mediated up-regulation of *Itgae* mRNA expression, despite not increasing *Spi1* mRNA levels (**Fig. 6c**). The critical requirement of PU.1 for the expression of the *Itgae* and *Itgb7* genes and the involvement of PU.1 in the TGF-β-induced increase in *Mcpt2* mRNA levels were still observed in peritoneal MCs (**Fig. 6d**).

**Fig. 6.**
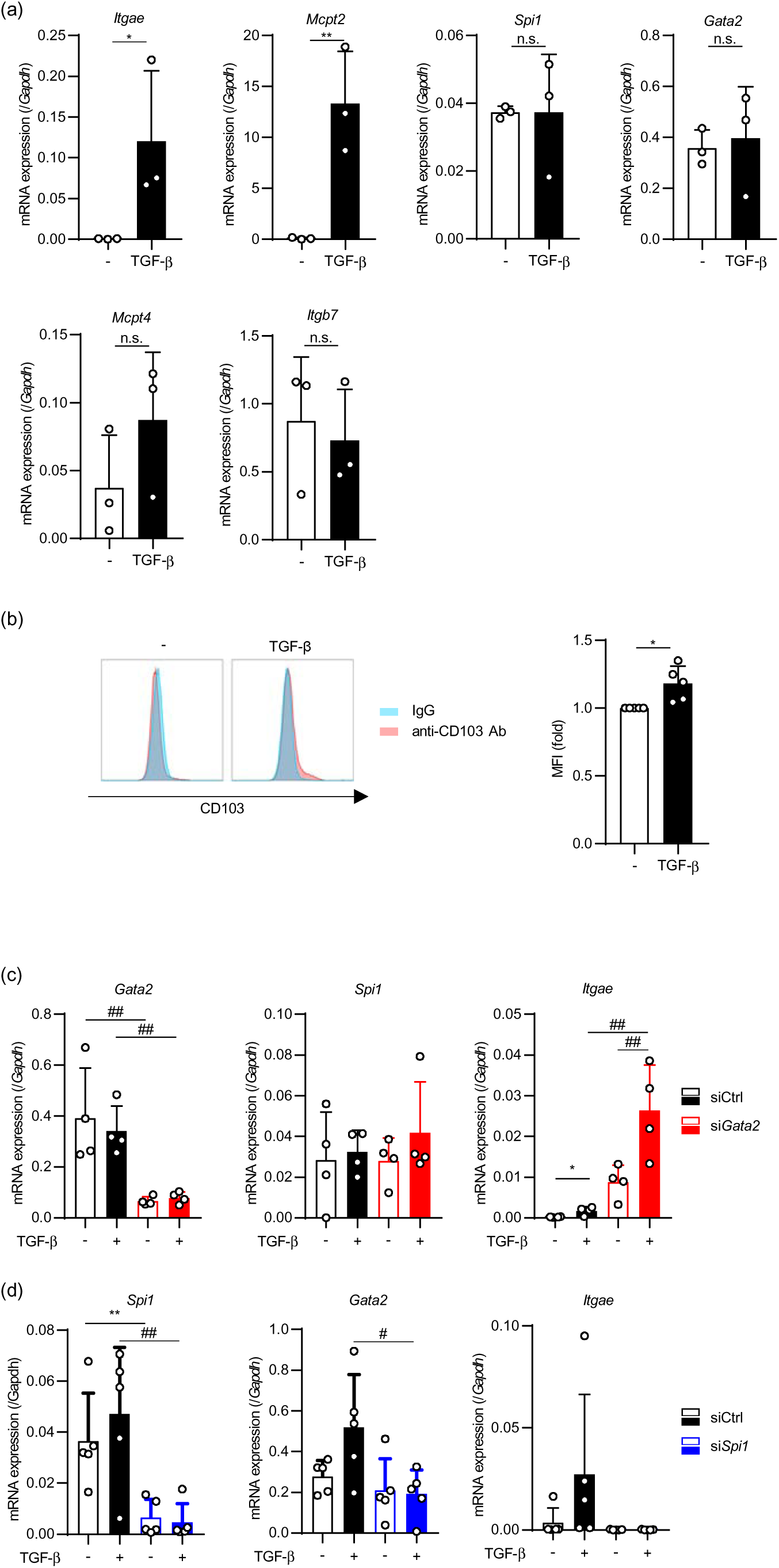
Roles of TGF-β signaling, GATA2, and PU.1 in CD103 expression on peritoneal MCs. **a.** and **b.** mRNA levels of MMC-related genes in (**a**) or surface CD103 on (**b**) peritoneal MCs incubated with or without TGF-β for 48 h. **c.** and **d.** Peritoneal MCs transfected with *Gata2* siRNA and its control (**c**) or with *Spi1* siRNA and its control (**d**) were stimulated with TGF-β for 48 h with same time schedule as those in Fig. 2e and 5e. Data represent the mean ± SEM of independent experiments. Dunnett’s multiple comparison test (**c**, **d**) and two-tailed paired *t*-test (**a**, **b**) were used for statistical analyses. *, *p* < 0.05; **, *p* < 0.01; n.s., not significant.

### PU.1 was involved in CD103 expression in DCs

Although PU.1 is a transcription factor that is essential for the development of DCs, the role of PU.1 in CD103 expression in DCs remains unclear. The ChIP-Atlas database suggested that region3, which we identified as a PU.1-binding site on the *Itgae* gene in MCs in the present study, was highly occupied by PU.1 in DCs (**Fig. 7a**). To evaluate the effects of *Spi1* KD on CD103 expression in DCs, Flt3L-induced BMDCs were generated. A flow cytometric analysis of BMDCs using specific markers for cDC and pDC showed that CD103 was mainly expressed in cDCs (**Fig. 7b**). The ratio of cDC/pDC was increased by the TGF-β stimulation and decreased by *Spi1* KD (**Fig. 7c**). The surface expression of CD103 on cDCs was markedly increased by the TGF-β stimulation, which was significantly suppressed by *Spi1* KD (**Fig. 7d**).

**Fig. 7.**
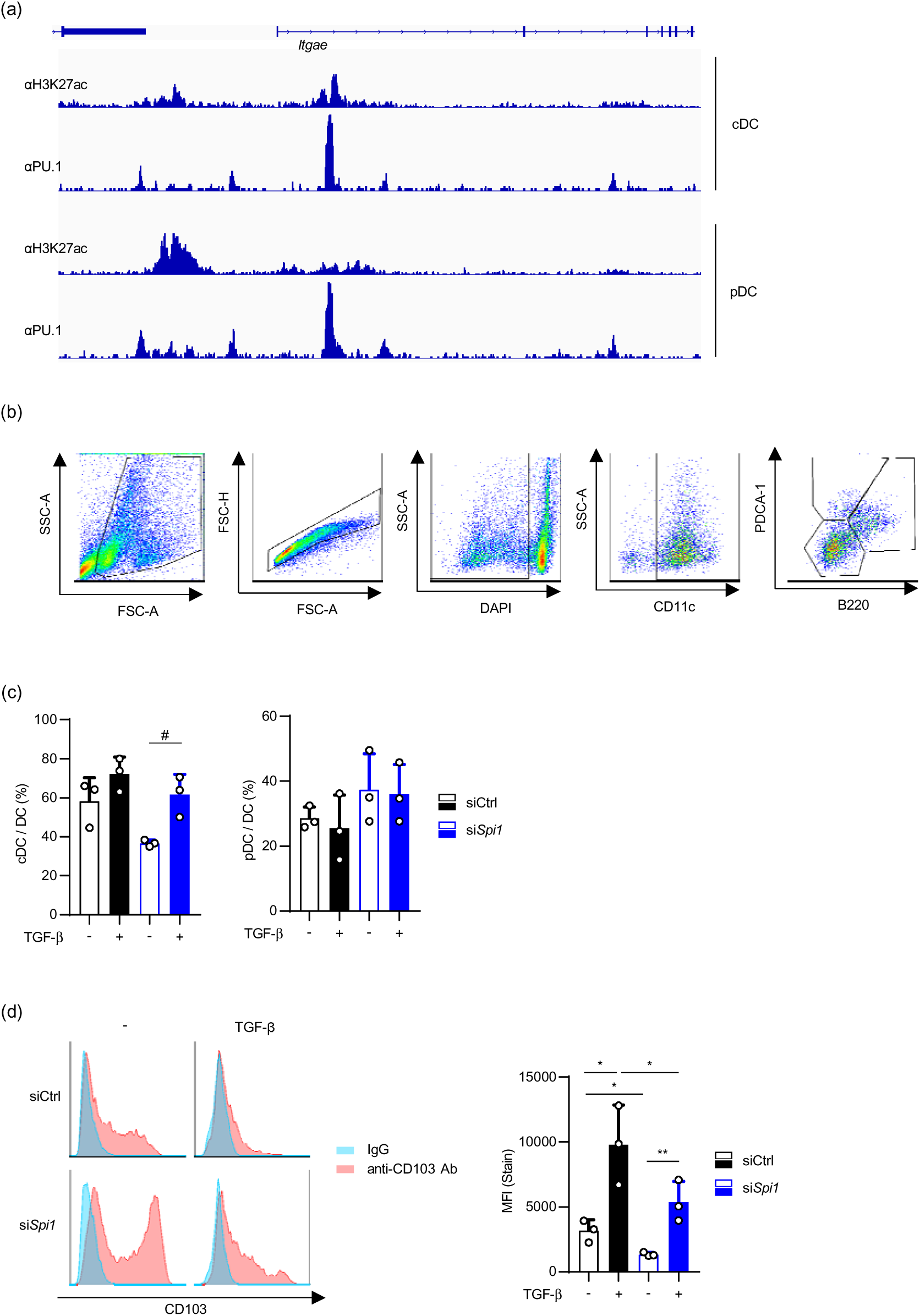
Roles of PU.1 and TGF-β-signaling in CD103 expression on DCs. **a.** ChIP profiles showing acetylation of H3K27 and PU.1 binding in the *Itgae* gene in DCs. H3K27 acetylation in cDC (https://chip-atlas.dbcls.jp/data/mm9/eachData/bw/SRX1255004), H3K27 acetylation in pDC (https://chip-atlas.dbcls.jp/data/mm9/eachData/bw/SRX1255006), PU.1-binding in cDC (https://chip-atlas.dbcls.jp/data/mm9/eachData/bw/SRX4909227), and PU.1-binding in pDC (https://chip-atlas.dbcls.jp/data/mm9/eachData/bw/SRX742829) were obtained via ChIP-Atlas. **b.** Gating strategy and dot-plot profiles showing surface CD103 levels on Flt3L-induced BMDCs. **c.** Ratio of cDC/pDC in TGF-β-treated and/or *Spi1* siRNA-transfected BMDCs. **d.** Effects of *Spi1* KD and the TGF-β stimulation on CD103 levels on cDCs in Flt3L-induced BMDCs. A typical histogram of CD103 on cDCs (left) and MFIs of CD103 on cDCs in three independent experiments (right). Data represent the mean ± SEM of independent experiments. Dunnett’s multiple comparison test (**c**, **d**) was used for statistical analyses. *, *p* < 0.05; n.s., not significant.

Collectively, these results indicate that PU.1 positively regulated the expression of CD103 on DCs.

## Discussion

MCs grow in BM, similar to other hematopoietic lineages, however, in contrast to other lineages, MCs are released from BM into peripheral blood at an immature stage. Immature MCs develop into mature stages, termed MMCs and CTMCs, in peripheral tissues in a manner that is dependent on environmental conditions. MMCs, CTMCs, and basophils, which highly and constitutively express FcεRI, play important roles in the IgE-dependent immune responses, including the onset of allergic diseases and the defenses against parasitic infections. In addition to these common characteristics, recent analyses of various gene targeted mice and human subjects have provided insights into the unique functions and gene expression profiles of each cell [19, 20]. To clarify the transcriptional mechanisms underlying cell-type specific gene expression, we previously examined transcription factors involved in the expression of the *Mcpt1* and *Mcpt2* genes and identified GATA2 and TGF-β signaling as positive regulators [1]. Although we observed a cooperative relationship between GATA2 and Smads in MMCs, it was insufficient to explain the commitment between MMCs and CTMCs because GATA2 is commonly required for the development and gene expression of MCs, basophils, and their common progenitors [21–23]. Therefore, in the present study, we investigated the regulatory mechanisms of CD103, another hallmark of MMCs, and found a novel role for GATA2 that negatively regulated surface CD103 and *Itgae* mRNA levels in MCs. The repressive effects of GATA2 on the *Itgae* gene were mediated through the regulation of PU.1. *Spi1* mRNA expression and PU.1 binding to the *Itgae* gene were significantly enhanced in *Gata2* KD MCs. In previous studies, the inhibitory role of GATA2 in PU.1-dependent gene expression was observed in various hematopoietic lineages, which are involved in the determination of hematopoietic cell fates [4, 9]. In contrast, PU.1 and GATA2 have been shown to cooperatively transactivate specific gene expression in MCs [10–12, 24]. The present results indicated that PU.1 was still repressed by GATA2 in MCs, even though PU.1 and GATA2 functioned synergistically in MCs. *Gata2* KD enhanced the expression of PU.1 and subsequently accelerated the recruitment of PU.1 to the *Itgae* gene. Further detailed analyses are needed to clarify whether PU.1 induces other MMC-related characteristics in MCs. In consideration of our previous studies showing that PU.1-overexpressing BMMCs acquired several monocyte-like characteristics [25–27], the TGF-β stimulation, which may enhance PU.1 functions, may induce the PU.1-depenent expression of other monocyte/MC-related genes.

The TGF-β-Smad axis induces the development and specific gene expression of MMCs [17, 28]. However, the transcriptional mechanisms underlying TGF-β-induced CD103 expression markedly differed from those responsible for the expression of Mcpt1/2, and were accompanied by the opposite roles of GATA2. In the case of the expression of Mcpt1 and 2, the complex of GATA2/Smad2/Smad4 transactivated the *Mcpt1* and *Mcpt2* genes via direct binding to the tandemly repeated *cis*-enhancing elements in these genes in TGF-β-stimulated MCs [1]. In contrast, the TGF-β-Smad axis supported the recruitment of PU.1 to the *Itgae* gene in MCs, which was inhibited by GATA2. We recently reported that TGF-β enhanced the development of basophils induced by C/EBPα and GATA2 [29]. These findings suggest that Smads activated in TGF-β-stimulated cells accelerate the recruitment and/or function of cell-type specific transcription factors that are involved in the terminal differentiation of each lineage.

Previous studies investigated the transcriptional regulation of CD103 in DCs and T cells. The involvement of IRF8 in CD103 expression on DCs was demonstrated using the CD11c-specific KO of *Irf8*, in which CD103^+^ migratory DCs were not detected in the lungs [30]. Furthermore, the DC-specific KO of Smad7, which was shown to accelerate the effects of the TGF-β stimulation in DCs, up-reulate the expression of IRF8 and Batf3 in DCs, and enhance the development of CD8^+^/CD103^+^ DCs in the spleen [31]. Another study reported a suppressive role for GATA2 in IRF8, with the lack of the *Gata2* enhancer up-regulating IRF8 expression in granulocyte-monocyte progenitors [32]. In addition, we previously found that MCs expressed IRF8 [33]. These findings prompted us to investigate the effects of *Irf8* KD on CD103 expression in MCs; however, *Itgae* mRNA levels were not decreased in *Irf8* KD BMMCs (data not shown). In the present study, *Irf4* KD did not reduce *Itgae* expression in MCs (**Fig. 3b**), despite IRF4 being required for the development of intestinal CD103^+^ DCs [18]. Based on these results, we conclude that PU.1 transactivated the *Itgae* gene in MCs in an IRF4- and IRF8-independent manner. Furthermore, this is the first study to show the essential role of PU.1 in CD103 expression in DCs, in the background of studies reporting the involvement of the PU.1-related transcription factors IRF4, IRF8, and Batf3, but not PU.1, in the expression of CD103 in DCs.

## Acknowledgments

We thank members of the Laboratory of Molecular Biology and Immunology, Department of Biological Science and Technology, Tokyo University of Science for their constructive discussions and technical support.

This work was supported by a Grants-in-Aid for Scientific Research (B) 23K26860 (CN), 23H02167 (CN), and 20H02939 (CN); a Grant-in-Aid for Early-Career Scientists 24K17872 (KN); a Research Fellowship for Young Scientists DC2 and a Grant-in-Aid for JSPS Fellows 21J12113 (KN); a Scholarship for Doctoral Students in Immunology (from the Japanese Society for Immunology to NI); a Tokyo University of Science Grant for President’s Research Promotion (CN); the Tojuro Iijima Foundation for Food Science and Technology (CN); a Research Grant from the Mishima Kaiun Memorial Foundation (CN); and a Research Grant from the Takeda Science Foundation (CN).

We greatly appreciate the consideration from Ms. Yayoi Yasuda, Dr. Masako Yasuda, and Dr. Kimihiko Yasuda.

## Authors’ contributions

K.I., and K.N. performed experiments, analyzed data, and prepared figures; Y.A., W.Z., M.I., and N.I. performed experiments; and analyzed data; K.K. designed the research, and performed experiments; C.N. designed the research, and wrote the manuscript.

## Supplemental information

None.

